# PUPAL COLOUR PLASTICITY AS A STRATEGY AGAINST DESICCATION

**DOI:** 10.64898/2026.05.18.725992

**Authors:** Bhanu Bhakta Sharma, Subhash Rajpurohit, Ullasa Kodandaramaiah

## Abstract

1. Terrestrial insects are vulnerable to desiccation due to their small body size. Because insects lose most water through cuticular evaporation, cuticular traits strongly influence desiccation tolerance. Individuals with greater cuticular melanisation, i.e., darker ones, are hypothesised to tolerate desiccation better than less melanised ones.
2. In many butterflies, pupal colour is plastic – individuals pupating on leaves tend to be greener, while those that pupate away from leaves (off-leaf), such as on tree bark or defoliated twigs, tend to be browner. Brown pupae are hypothesised to have more cuticular melanin and are expected to experience higher desiccation stress than leaf-borne green pupae. Thus, plasticity in pupal melanisation may be an adaptation against desiccation. We tested this in the butterfly *Eurema blanda*.
3. We demonstrate that individuals pupating on on-leaf substrates are greener than those pupating on off-leaf substrates, and that desiccation stress is higher in the off-leaf substrates, a microenvironment typical of brown pupae, than in typical green pupae. Using Raman spectroscopy, we show that brown, but not green, pupal cuticles contain melanin.
4. Following this, we obtained greener and browner pupae by manipulating substrate colour. When subjected to desiccation stress, browner pupae survived better than greener ones. There was no correlation between pupal colour and survival in the absence of desiccation stress. Thus, melanisation appears to confer a survival advantage to pupae by increasing desiccation tolerance.
5. Survival under desiccating conditions was inversely related to water loss. Interestingly, melanisation did not correlate with water loss, suggesting that melanisation helps tolerate desiccation through physiological mechanisms not directly related to water loss reduction.
6. Our findings reveal an additional, crucial, adaptive value of pupal colour plasticity, a trait that has been studied primarily from an anti-predatory perspective.

## INTRODUCTION

Water is essential for all animals, and there are numerous adaptations across the animal kingdom for water intake from the external environment and to prevent water loss. Insects are particularly vulnerable to water loss because they are small and have a large surface area to volume ratio, which creates potential for rapid water loss through cuticular transpiration (Hadley, 1994; Kühsel et al., 2017; O’Donnell, 2022; Rajpurohit et al., 2016). Indeed, cuticular transpiration accounts for the highest proportion of total water loss in insects (Gibbs & Rajpurohit, 2010; Kühsel et al., 2017; Rourke, 2000) with one study on the harvester ant *Pogonomyrmex barbatus* reporting 97% water loss through this route (Johnson & Gibbs, 2004). The primary abiotic factor influencing the rate of cuticular water loss is the difference in vapour pressure between the free water within the organism and the water in the surrounding environment, i.e., vapour pressure deficit (VPD). VPD increases with increase in temperature and decreases with relative humidity (RH) (Howard et al., 2020; Kleynhans & Terblanche, 2011; O’Donnell, 2022).

Despite being intrinsically prone to high rates of cuticular water loss, insects are among the most successful groups, and have radiated across diverse habitats ranging from humid to arid. Among the factors underlying their evolutionary success are their adaptations for survival in highly desiccating environments (Benoit et al., 2023; O’Donnell, 2022; Sinclair et al., 2024). Not surprisingly, many adaptations against desiccation predominantly involve cuticular traits. The outermost cuticular layer, i.e., epicuticle, contains hydrophobic compounds such as hydrocarbons and melanin, which limit water loss through transpiration (King & Sinclair, 2015; Ramniwas et al., 2013). A prominent cuticular property thought to vary in relation to desiccation stress levels is melanisation. There is intraspecific variation in cuticular melanisation both within populations (King & Sinclair, 2015; Parkash, Rajpurohit, et al., 2008; Parkash, Sharma, et al., 2009; Parkash, Singh, et al., 2009; Rajpurohit et al., 2008) and between populations distributed in disparate habitats (Parkash, Sharma, et al., 2009; Rajpurohit et al., 2008). The rate of cuticular water loss is significantly less for melanised (darker) individuals compared to non-melanised (lighter) ones (Farnesi et al., 2017; King & Sinclair, 2015; Parkash, Sharma, et al., 2008; Ramniwas et al., 2013; but see Matute & Harris, 2013; Rajpurohit et al., 2016; Wittkopp et al., 2011). The majority of these studies have been on *Drosophila* (e.g., Hoffmann & Harshman, 1999; Kellermann et al., 2018). For instance, there is a negative correlation between cuticular melanisation and the rate of water loss at both the intrapopulation and interpopulation levels, in wild and lab-reared populations of *Drosophila melanogaster* (Parkash, Sharma, et al., 2009).

In holometabolous insects, life stages can vary strongly in mobility, and, therefore, in their adaptations against desiccation. Feeding stages such as larvae and adults can include various behavioural mechanisms, including drinking water to increase water intake, obtaining water from the diet to compensate for water loss, and moving to moist microhabitats to avoid evaporative water loss (Chown, 2002; Guedes et al., 2015; O’Donnell, 2022). On the other hand, eggs and pupa are likely more vulnerable to desiccation stress because they are immobile, and cannot rely on such behavioural mechanisms (discussed in Klockmann & Fischer, 2017).

Many butterflies exhibit plasticity in pupal colour, ranging from brown to green (Hiraga, 2006; Maisch & Bückmann, 1987; Mayekar & Kodandaramaiah, 2017; Piñones-Tapia et al., 2017; Smith et al., 1988; Yamamoto et al., 2011; Yumnam et al., 2021). In many butterflies pupal colour is determined by pigments deposited on the epicuticle (Ohtaki & Ohnishi, 1967; Yoda et al., 2020). Brown pupae are thought to have melanin while green pupae have blue and yellow pigments (Maisch & Bückmann, 1987; Ohnishi, 1959; Ohtaki & Ohnishi, 1967; Phelps et al., 2024; Yoda et al., 2020). Pupal colour tends to be correlated with pupation substrate – green pupae are formed primarily under leaves (hereafter, on-leaf) substrates, and brown pupae on either substrates away from leaves (hereafter, off-leaf) such as soil, stem and rocks, or under defoliated leaf midribs (Clarke & Sheppard, 1972; Mayekar & Kodandaramaiah, 2017; Ohnishi, 1959; Yumnam et al., 2021). Pupal colour plasticity is widely thought to have evolved as a strategy against predation (Hazel et al., 1987; Lindstedt et al., 2019; Piñones-Tapia et al., 2017; Stefanescu, 2004), because green and brown pupae are presumably better camouflaged when viewed against leaf and off-leaf substrates respectively (reviewed in Mayekar & Kodandaramaiah, 2017; Piñones-Tapia et al., 2017).

Pupation substrates may also differ in their microclimatic variables, and, hence, the extent of desiccation stress experienced by pupae. Compared to brown ones, green pupae may experience weaker desiccation stress due to i) transpiration by leaves, which releases moisture, and ii) their sheltered position under the leaves, reducing direct exposure to sunlight. Thus, the brown pupal phenotype may have evolved as an adaptation against desiccation stress (Mayekar & Kodandaramaiah, 2017). In the butterfly *Mycalesis mineus*, a higher proportion of brown pupae develop when reared under low RH conditions than at high RH (Mayekar & Kodandaramaiah, 2017), providing indirect support for this idea. However, whether pupal colour influences desiccation tolerance has not been experimentally tested.

We here tested the melanin desiccation hypothesis in relation to pupal colour plasticity in the butterfly *Eurema blanda* (Pieridae: Pierinae). Our laboratory observations suggested that *E. blanda* shows pupal colour plasticity as a continuum, ranging from brown to green, with intermediates. We quantify pupal colour and show in experiments that pupae forming on leaves are predominantly green, whereas those on off-leaf substrates are brown. We established using Raman spectroscopy that cuticles of brown, but not green, pupae contain melanin. We used microclimatic data to test whether VPD – a measure of desiccation stress differs between the typical substrates of green and brown pupae. We predicted that, under strongly desiccating conditions:

i. the degree of melanisation is inversely correlated with the rate of water loss (Prediction 1)
ii. survival is inversely correlated with the rate of water loss (Prediction 2)
iii. survival is positively correlated with the degree of melanisation (Prediction 3).

Melanin may confer a survival advantage even in the absence of desiccation stress, for instance, by increasing immunity (reviewed by Dubovskiy et al., 2013; Fedorka et al., 2013). Thus, darker pupae may generally survive better than lighter ones, irrespective of the presence of desiccation stress. On the other hand, if melanin functions primarily as an adaptation against desiccation, darker pupae should have a survival advantage in desiccating but not in non-desiccating conditions. We assessed the benefit of melanin under both desiccating and non-desiccating conditions. We predicted that (iv) there is no relationship between survival and degree of melanisation under non-desiccating conditions (Prediction 4).

## MATERIALS AND METHODS

### Experimental overview

Pupal colour was quantified to test for differences between pupae on leaves and off-leaf substrates (Experiment 1). Raman spectroscopy was used to assay the presence of melanin on green and brown pupal cuticles (Experiment 2). Temperature and RH data from leaf and off-leaf substrates were used to test whether VPD differs between the substrates (Experiment 3). Two independent experiments were conducted to test the effect of pupal colour on desiccation tolerance. These experiments differed in the whether desiccation was imposed or not. Pupae were subjected to desiccation in experiment 4 , but not in experiment 5, which allowed us to test whether melanisation is advantageous under both desiccating and non-desiccating conditions (Prediction 3 and 4).

### Experiment 1: Pupation substrate in relation to pupal colour

*E. blanda* in the study locality uses *Cassia fistula* as its host plant (personal observation). *E. blanda* larvae were reared under outdoor conditions in sleeves made of nylon mesh. Each sleeve enclosed the larvae and twigs with tender leaves. Late instarlarvae were collected and, brought to the laboratory and reared further in metal cages (40 × 40 × 40 cm; solid aluminium frame fitted with ∼49-holes/cm² mesh) until pupation. *C. fistula* twigs dipped in water served as larval food and potential pupation substrates. The twigs were placed in a way that allowed wandering larvae (late-instar larvae prior to pupation) to access all interior areas of the rearing cages and potentially pupate anywhere. On the second day after pupation, the pupation substrate was recorded as on-leaf (when pupation formed on leaves) or off-leaf (when pupae formed on stems or artificial substrates), and pupal colour was quantified (see below ‘Quantification of pupal colour’).

### Experiment 2: Melanin assay

Lateral cuticular sections of dry pupal cases (remaining after successful butterfly eclosion) from green and brown pupae reared in independent experiments were fixed on glass slides. Raman spectra of these sections were obtained using a Horiba Xplora Plus Confocal Micro Raman Microscope (Horiba Jobin, France). Samples were excited with a 532 nm at 0.1 and 10 mW laser power settings, and Raman shift values between 1000 and 1800 cm⁻¹ were recorded. Synthetic melanin (Sigma-Aldrich Chemical Corporation, M0418) was used as a reference. A more detailed experimental protocol is available in Supp. Mat. Text S2.

### Experiment 3: Vapour pressure deficit assay

Dataloggers (Elitech, RC-4HC) were used to record temperature and RH at leaf and off-leaf substrates on wild *Cassia fistula* plants, representing the typical microhabitats of green and brown pupae. Data were collected from 18 loggers on each microhabitat type, and recorded every minute from from 08:30 to 17:00 h for 3 days. VPD was calculated using temperature and RH values. A more detailed experimental protocol is available in Supp. Mat. Text S3.

### Butterfly rearing for Experiments -5

*E. blanda* eggs were collected from wild *Cassia fistula* plants, and kept in a growth chamber (PLE-105; Pooja Lab Equipments, Mumbai, India) set at 27°C and 85% RH. A 12:12 hr dark:light cycle was maintained in all experiments. Hatchlings were provided *ad libitum* fresh leaves of *C. fistula*. Larvae that reached the wandering phase were carefully hand-picked and sexed by examining the presence or absence of testes located in the dorsal region of the abdominal segments (Retnakaran, 1970). Green and brown pupae were obtained by placing the pre-sexed wandering larvae in a paper cylinder (fitted with a perforated petri-plate at both ends) prepared from green and black paper, respectively (Supp. Mat. Fig. S1). On the second day of pupation, pupal colour was quantified (details in ‘Quantification of colour in Supp. Mat. Text S1), and pupal weight was recorded using a sensitive weighing balance (MSA6.6S–000–DM 29405385, Sartorius Weighing Technology GmbH, Goettingen, Germany) with a precision of 1 µg. Henceforth, initial weight was considered a proxy of body size (Bauerfeind & Fischer, 2007).

### Experiment 4: Water loss and survival under desiccation stress

After being photographed and weighed, pupae were placed on a perforated shelf inside a transparent glass desiccator. The desiccator was maintained at <10 % RH, 25°C, by placing silica gel (Sigma-Aldrich, 13767) on the bottom of the desiccator. The silica gel was dried overnight at 60°C. Data loggers (HTC PDF LOG, Hazari Tech Connect, Mumbai) were used to monitor the RH and temperature inside the desiccator. Pupae were subjected to desiccation stress for 72 hours, which corresponds to approximately half the entire pupal stage (5 -7 days). Pupae were weighed immediately afterwards. Weight loss during the desiccation assay was assumed to be water loss (Bujan et al., 2016; Parkash, Sharma, et al., 2009; Rajpurohit et al., 2016). Hence, weight loss is henceforth referred to as water loss. Water loss during the desiccation assay was calculated gravimetrically (Benk et al., 2020; King & Sinclair, 2015; Kühsel et al., 2017) using the equation (adopted from Benk et al., 2020)

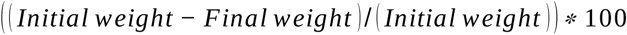

Here, initial and final weight refer to pupal weight before and after desiccation exposure.

The desiccation treated pupae were placed in 100ml pre-labeled Tarson tubes covered with a nylon mesh instead of the lid, at 25°C and 85% RH, and eclosion success was recorded. Butterflies were considered to have eclosed successfully only if all body parts were detached from the pupal case. Butterflies were considered to have survived the desiccation stress if they eclosed (Hayward et al., 2004; Klockmann & Fischer, 2017).

### Experiment 5:Survival in the absence of desiccation stress

The steps involved in rearing larvae and obtaining pupae were similar to that in experiment 4 with larvae originating from independently collected eggs from the wild population. The difference here was that pupae were not exposed to desiccation stress, and the pupal weight was measured only once on the second day after pupation. Briefly, the eclosion success of photographed and weighed pupae was recorded by keeping the pupae on the prelabeled 100ml Tarson tube covered with nylon mess in 27C, 85% condition.

### Experiment 6 and 7

We conducted additional experiments following protocols similar to that of Experiments 4 and 5, with the larvae originating from independently collected eggs from the wild. The larvae were reared and pupated at 60% RH. Thus, Experiments 6 and 7 tested the effect of pupal lightness on survival under desiccated and non-desiccated conditions, when grown at different RH. These additional experiments were done to test whether the relationship between pupal colour and desiccation tolerance differs depending on the humidity experienced during early development.

### Quantification of pupal colour

The lightness of pupae was assumed to be inversely correlated with the amount of melanin pigmentation, in line with established protocols from other studies that have quantified melanin from biological samples (Britton & Davidowitz, 2024; Farnesi et al., 2017; Gautam & Kunte, 2020; Hegna et al., 2013; Kutama et al., 2024; Lopez et al., 2021; Pool & Aquadro, 2007; Ramniwas et al., 2013). Photographs of the pupae were taken inside a white custom-built thermocol light box (lbh: 29cm x 29cm x 49cm), two opposite sides of which were fitted with 2 LED strips (4V) having 36 diodes, as a constant source of light. Each pupa was placed on a standard gray card (18% gray) flanked by a white and black piece of gray (JJC-white-balance-gray-GC-2), a scale in mm, and an individual ID for each pupa and photographed. All pupae used in the analysis were photographed with a Nikon D3200 Digital SLR camera (Nikon, Tokyo, Japan) and a Sigma 105 mm macro lens (Sigma Corporation, Kanagawa, Japan). Exposure correction of pictures was done using ImageJ 1.53t (Schneider et al., 2012). The pupal area was extracted using the Paint 3D app v. 6.20003.4017.0 (<u>Paint 3D–Microsoft Store Apps</u>). A custom-written macro script was run in ImageJ to extract the pixel value average (0 - 255) separately for red, green, and blue. The lightness of each photo was calculated as ((R+G+B+)/(3)) (Roberts et al., 2022). R, G, and B channel values for each photo are mean values of Red, Green, and Blue, respectively. Greenness Index (GI), an estimate of how green a pupa is, with higher and lower values representing greener and browner pupae respectively (Yumnam et al., 2021), was calculated as ((G*2)/(B+R)).

### Statistical analyses

Data analysis and visualisation were done in R v. 4.5.2 (R Core Team, 2024) via R Studio v. 2025.9.2.418 ‘Cucumberleaf Sunflower’ (Posit team, 2024). The frequency distribution of GI of pupae from all experiments was plotted to visualize variation in colour. Because Shapiro-Wilk tests indicated that GI and lightness were not normally distributed, a Spearman’s correlation test, using the function ‘cor.test’, was used to test the correlation between the two variables. A Wilcoxon rank-sum test was done using the function ‘wilcox.test’ to test whether GI differed between leaf and off-leaf pupae in Experiment 1. Similarity between each pupal Raman spectrum and the reference spectrum was quantified across the full spectral range using Pearson correlation coefficients calculated with the ‘cor’ function on baseline corrected (using asymmetric least squares method) and maximum normalized spectra (Molina et al., 2024; Mostafapour et al., 2026; Samuel et al., 2021). For pupae scanned at 10 mW, two spectra per individual were recorded, while pupae scanned at 0.1 mW had only a single spectrum per individual for analysis. Spectra belonging to the same pupa were averaged to obtain a single mean spectrum. A Pearson correlation coefficient was then calculated between the mean pupal spectrum and the reference spectrum. These pupal correlation coefficients were grouped by pupal colour (brown or green) and mean correlation values were reported for each group. A Linear Mixed-effects Model (LMM) was used to test the difference in VPD between on-leaf and off-leaf microhabitats. Mean VPD values were calculated for each time point and microhabitat type. Model fitting was done using the *lmer* function in the *lmerTest* package v. 3.1.3 (Kuznetsova et al., 2017) To account for repeated measures, we tested the effect of the main predictor variable, substrate (on-leaf and off-leaf), on VPD, including day, time of day, and datalogger ID as random intercepts.

For Experiments 4-7 data on survival were represented as 1 (survived) and 0 (dead). GLMs (Generalised Linear Models) were built with the ‘glm’ function, assuming a binomial family distribution and the default *‘logit’* link to test the effect of lightness and sex on survival. Both the additive models (Lightness + Sex) and an interaction models (Lightness * Sex) were built. Model selection was based on Akaike Information Criterion (AIC), and the model with less AIC was retained (Palombo et al., 2020; reviewed in Sutherland et al., 2023). The pathways linking lightness to survival was estimated using structural equation models (SEM) (Britton & Davidowitz, 2024; Debecker et al., 2015) using the ‘sem’ function from the *lavaan* package v. 0.6.21 (Rosseel et al., 2012) for data from Experiments 4 and 5. SEM allows the simultaneous estimation of direct and indirect effects of a predictor on a response variable via multiple parallel mediators (Gunzler et al., 2013). Initial body weight and water loss were included as potential parallel mediators, while sex was included as a covariate in Experiment 4. No causal order between the mediators was assumed. Because survival was binary, the model was estimated using Weighted Least Squares Mean and Variance adjusted (WLSMV) estimator with a default *‘probit’* link (reviewed in Newsom & Smith, 2020; Rosseel et al., 2012). SEM based mediation analysis for Experiment 5 was conducted similar to Experiment 4, except that only initial body weight was included as a potential mediator and not water loss, because it was not quantified. Standardized estimates from the models were reported. SEM model fit was assessed using the Comparative Fit Index (CFI), Tucker–Lewis Index (TLI), and Root Mean Square Error of Approximation (RMSEA), with values of CFI and TLI ≥ 0.95 and RMSEA ≤ 0.05 indicating good model fit (reviewed in Gunzler et al., 2013).

## RESULTS

GI and lightness of *E. blanda* pupae were bimodally distributed, with the peaks representing green and brown pupae (Supp. Mat. Fig S2). Across all experiments, GI was strongly positively correlated with lightness, such that greener pupae were lighter (Supp. Mat. Fig. S4).

### Experiment 1: Pupation substrate in relation to pupal colour

The GI of pupae on leaf substrates (median = 1.39; n = 39) was significantly higher than that of pupae on off-leaf substrates (median = 1.180; n = 52) (Wilcoxon rank-sum test; *W* 244, p-value < 0.001; Fig. 1).

**Figure 1:**
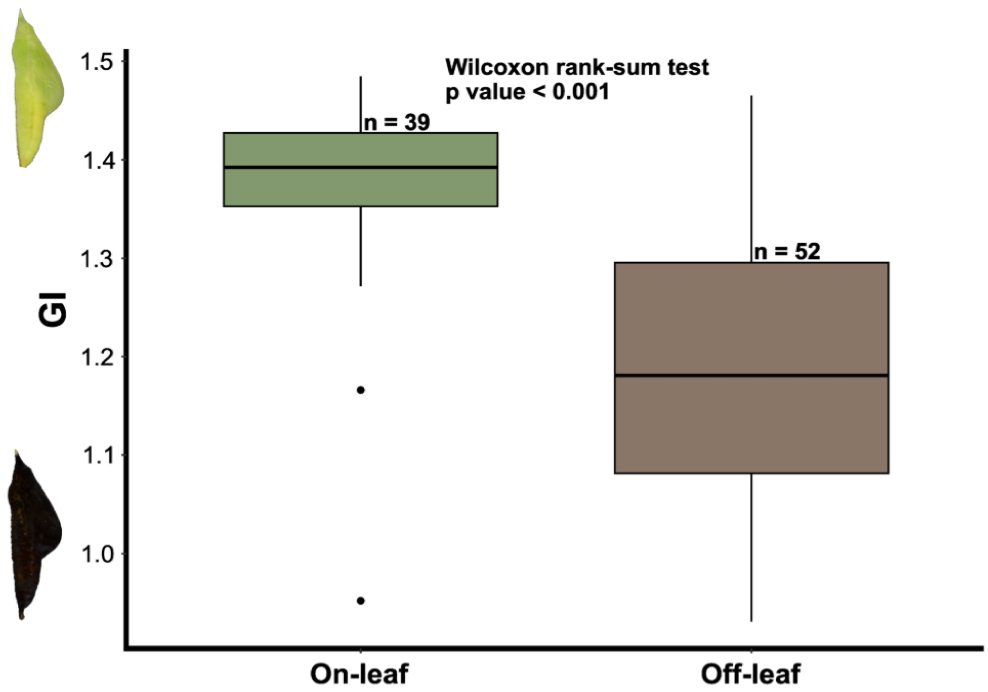
Comparison of pupal greenness index (GI) between on-leaf and off-leaf pupation substrates. Box plots show the median (horizontal line), interquartile range (box), and range excluding outliers (whiskers), with outliers plotted as individual points. Numbers above boxes indicate sample sizes. Representative images of green and brown pupae are shown alongside the y-axis.

### Experiment 2: Melanin assay

The average GI of the brown pupae used for the Raman intensity measurements was 1.034 (± 0.043 SD; n = 8), and that of the green pupae was 1.391 (± 0.055 SD; n = 8). These values correspond to the peaks of green and brown pupae (Supp. Mat. Fig. S2). Brown pupae had two peaks in their Raman spectra – a strong one about 1585 cm⁻¹ and a smaller one at about 1385 cm⁻¹ - at both 0.1 mW and 10 mW laser powers. Green pupae had multiple small peaks (Supp. Mat. Fig. S6). Raman spectral similarity to the reference spectrum differed between pupal morphs. Pearson correlation coefficients showed that brown pupae exhibited higher similarity to reference than green pupae at both laser power settings. Pupal spectrum obtained at 10mW, when compared with the reference obtained at 10mW, yield higher correlation coefficient for brown (n = 3, mean = 0.77, SD ± 0.014) than for green pupae (n = 3, mean 0.343, SD ± 0.0124). A similar pattern was observed when pupal spectrum were compared with reference obtained at 0.1 mW with brown pupae again showing higher similarity (n = 3, mean 0.407, SD ± 0.006) than green pupae (n = 3, mean 0.184, SD± 0.006). Likewise, pupal spectrum obtained at 0.1mW, when compared with the reference obtained at 10mW, yield higher correlation coefficient for brown (n = 6, mean = 0.545, SD ± 0.184) than for green pupae (n = 6, mean 0.32 SD ± 0.057). A same trend was evident when pupal spectrum were compared with reference obtained at 0.1 mW with brown pupae again showing higher similarity (n = 6, mean 0.315, SD ± 0.113) than green pupae (n = 6; mean 0.196, SD± 0.027).

### Experiment 3: Vapour pressure deficit assay

VPD was significantly higher in off-leaf than in leaf substrates i.e. off-leaf substrates were drier than leaf substrates (LMM estimate = 0.715, SE± 0.086 p < 0.001, Supp. Mat. Fig. S5). The model was based on 17,280 recordings across 510 time points.

### Experiment 4: Water loss and survival under desiccation stress

The analysis was conducted on 169 pupae. In the additive model, with lightness as a continuous predictor and sex as categorical predictor the survival probability of pupae under desiccation stress was negatively affected by pupal lightness (GLM estimate = − 0.02, z = −2.929, p = 0.003, Fig. 2A). Survival did not differ between males and females (GLM estimate = −0.11, z = −0.306, p = 0.76). In the interaction model, adding a Lightness X Sex interaction did not improve model fit (AIC = 175.95 vs 174.08 for the additive model ΔAIC = 1.87), and the interaction was not significant (GLM estimate = − 0.004, z = −0.357, p = 0.72, Supp. Mat Table S1a). We therefore retained the additive model.

**Figure 2A:**
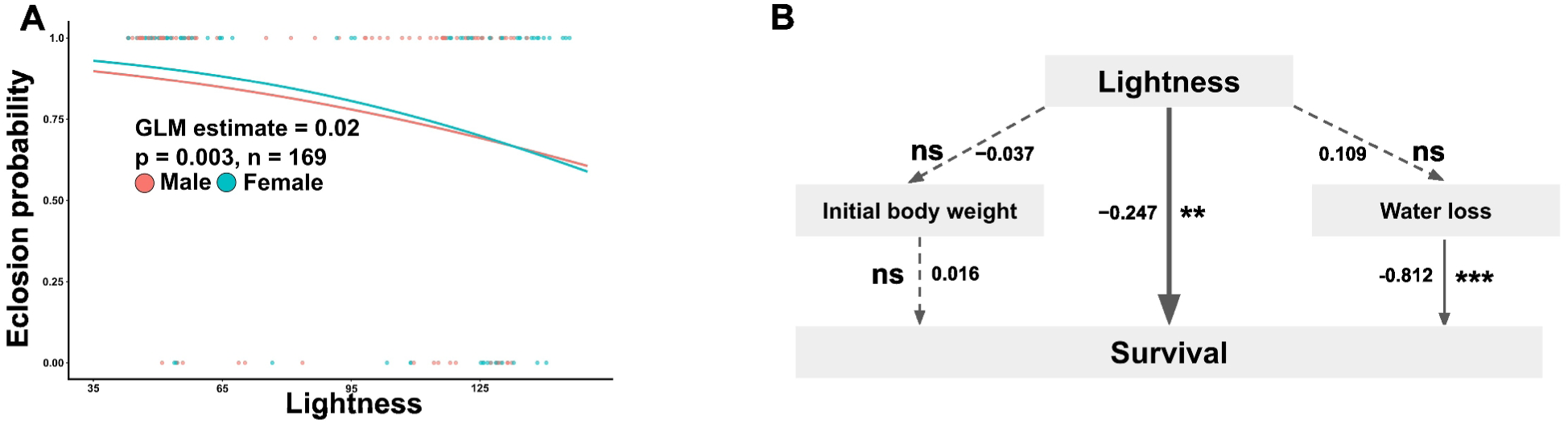
Effect of pupal lightness on eclosion probability of pupae under desiccation stress. The dots represent data points, and regression curves for each sex are shown as solid lines. Estimates are for the independent effect of lightness on eclosion probability. The statistical results shown on the graph correspond to the combined data for males and females. **Figure 2B: Path analysis depicting the direct and indirect pathways linking lightness to survival via mediators.** Numbers along paths indicate model estimates. Dashed arrows indicate indirect pathways while solid arrow indicate direct pathway. Labels “ns” indicate non-significant effects, while asterisks denote levels of statistical significance (*p = 0.01-0.05, **p = 0.001-0.01, ***p < 0.001)

In the SEM based mediation analysis of survival, lightness did not predict either initial body weight (estimate = −0.037, z = −0.445, p = 0.656) or water loss (estimate = 0.109, z = 1.112, p = 0.266). Similarly, sex did not predict either initial body weight (estimate = 0.013, z = 0.173, p = 0.863) or water loss (estimate = 0.009, z = 0.100, p = 0.920). Water loss predicted survival, with higher water loss reducing survival probability (estimate = −0.812, z = −21.647, p < 0.001). Initial body weight did not predict survival (estimate = 0.016, z = 0.160, p = 0.873). The indirect effect of lightness on survival via body weight was not significant (estimate −0.001, z = −0.150, p = 0.880). Similarly, the indirect effect via water loss was not significant (estimate = −0.089, z = −1.102, p = 0.270). The total indirect effect of lightness on survival via the two mediator was not significant (estimate = −0.089, z = −1.109, p = 0.268). The total effect of lightness on survival (estimate = −0.336, z = −2.846, p = 0.004, Fig, 2B) closely matched the direct effect (estimate = −0.247, z = −2.844, p = 0.004), indicating that the relationship between lightness and survival was direct and not mediated by body weight or water loss (Fig. 2B) (Gunzler et al., 2013).

### Experiment 5: Survival in the absence of desiccation stress

The analysis was conducted on 139 pupae. In the additive model, with lightness as a continuous predictor and sex as categorical predictor the survival probability of pupae in the absence of desiccation stress was not affected by pupal lightness (GLM estimate = −1.776, z = 0.892, p = 0.37, Fig. 3A). Survival did not differ between males and females (GLM estimate = −22.193, z = −0.003, p 0.998). In the interaction model, adding a Lightness X Sex interaction did not improve model fit (AIC = 11.56 vs 9.55 for the additive model; ΔAIC = 2.01), and the interaction was not significant (GLM estimate = −1.78, z = −0.007, p = 0.995, Supp. Mat. Table S1a). We therefore retained the additive model. In the SEM based mediation analysis of survival, the indirect effect of lightness on survival via body weight was not significant (estimate = 0.002, z = −0.861, p = 0.389). The total effect of lightness on survival was not significant (estimate = −0.627, z = −0.318, p = 0.751) indicating that lightness did not influenced survival in the absence of desiccation stress.

**Figure 3:**
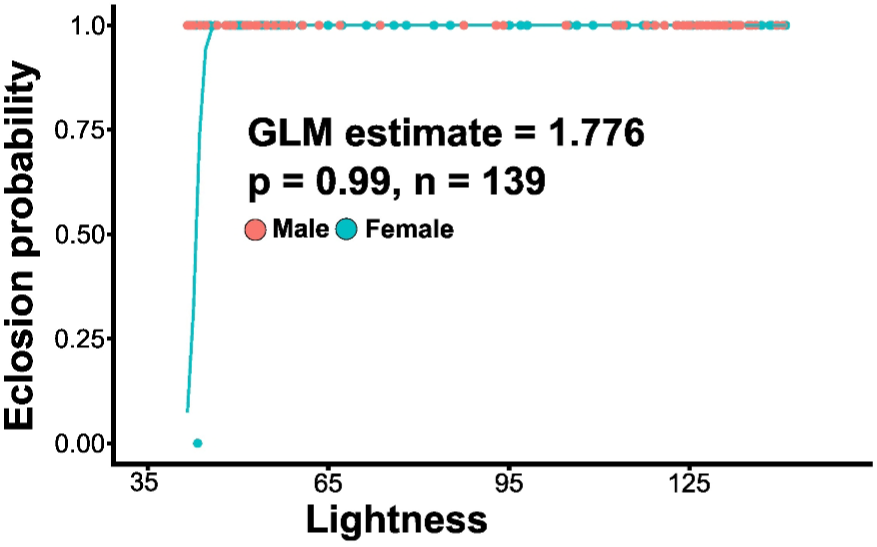
Effect of pupal lightness on eclosion probability of pupae in the absence of desiccation stress. The dots represent data points, and regression curves for each sex are shown as solid lines. Estimates are for the independent effect of lightness on eclosion probability in the absence of desiccation stress. The statistical results shown on the graph correspond to the combined data for males and females.

Results of Experiments 6 and 7 (relationship between pupal lightness and survival) are detailed in Supp. Mat. Table S1a. In brief, survival was correlated with lightness (GLM estimate = −0.01, z =−2.38 p = 0.017, n = 115; Supp. Mat. Fig. S7A) in Experiment 6, i.e., under desiccation stress, but there was no relationship between the two (GLM estimate = −0.01, z = −2.38 p = 0.79, n = 138; Supp. Mat. Fig. S7B) in Experiment 7, i.e., in the absence of desiccation stress.

## DISCUSSION

Results from Experiment 1 confirm that pupae on off-leaf substrates tend to be brown while those on leaves tend to be green (Fig. 1). Although brown pupal colour has widely been assumed to be due to melanin, to our knowledge, ours is the first study to biochemically characterize the presence of melanin in the cuticle of brown pupae using Raman spectroscopy. The broad Raman peaks near 1385 and 1585 cm⁻¹ have been consistently detected in synthetic and natural melanins across multiple excitation wavelengths in other studies (e.g., Galván et al., 2013; Huang et al., 2004). These peaks correspond to the characteristic D and G bands typically observed in the Raman spectra of disordered graphite (reviewed in Galván et al., 2013; Huang et al., 2004). The G band (∼1585 cm⁻¹) arises from the stretching vibrations of hexagonally arranged carbon rings, while the D band (∼1385 cm⁻¹) is attributed to linear stretching of three of the six C-C bonds. The Raman spectral similarity to the reference spectrum differed between pupal morphs, with brown pupae showing higher similarity than green pupae (Khan & Madden, 2012; Samuel et al., 2021; Willemse-Erix et al., 2009). This further corroborates the presence of melanin in brown pupae but not in green pupae.

Results from Experiment 3 confirm that vapour pressure deficit is higher on off-leaf substrates than under leaves (Supp. Mat. Fig. S5). Thus, desiccation stress is higher when pupation happens away from leaves. Experiment 4 tested the relationship between pupal survival, colour and water loss. The survival of pupae was strongly affected by the amount of water lost during the desiccation assay – individuals losing more water had a lower survival (Fig. 2B). Lightness, a measure of melanisation, influenced survival under desiccating conditions, darker individuals, i.e. those that presumably invested more in melanin deposition on the cuticle had greater tolerance to desiccation. However, melanisation did not provide any survival advantage under non-desiccating conditions (Fig. 3). Our prediction that darker pupae will have lesser water loss was not supported. Our results highlight the complexity of the relationship between melanisation, water retention, and survival under desiccating conditions.

Pupal colour plasticity has traditionally been studied from an anti-predatory perspective (Hazel et al., 1987; Lindstedt et al., 2019; Piñones-Tapia et al., 2017), and primarily in temperate butterflies which may not experience strong desiccation stress. Our results indicate that melanisation serves a broader function, contributing not only to predator avoidance but also to tolerate desiccating conditions.

### Importance of water conservation

In Experiment 4, one-day-old pupae by which time the cuticle had already hardened – were subjected to desiccation stress for roughly 50% of their total pupal duration. After this period, they were returned to humid conditions, and eclosion success was monitored. We found a negative correlation between water loss under desiccating conditions and survival as measured by the probability of successful eclosion. During the pupal phase, the body of insects undergoes complete reorganisation – the larval structure breaks down and is restructured to form the adult anatomy (Gilbert, 2009). This reorganisation involves various complex processes, and water is essential to ensure these transformations occur normally. Our results suggest that water loss during a substantial portion of the pupal stage beginning around 24 hours after pupation, when the cuticle has already hardened may cause irreversible physiological damage. Given their immobility and limited ability to actively intake water from the environment (Yoder & Denlinger, 1991), pupae in this initial window may be especially vulnerable to desiccation.

One of the major factors influencing water loss is body weight (Kühsel et al., 2017). In insects, water loss tends to be negatively correlated with body weight (Guedes et al., 2015; Lighton et al., 1994). A larger body size translates to lower surface area:volume ratio, reducing the water loss rate (Howard et al., 2020; Kühsel et al., 2017). However, this pattern was not evident in Experiment 4, suggesting that body weight alone may not fully explain variation in water loss. Other physiological processes, such as spiracular respiratory water loss and the involvement of protective metabolites may play a role in regulating water loss.

### Melanin pigmentation is adaptive under desiccating conditions

Both lightness and GI influenced eclosion success under desiccating conditions (Fig. 2A & C for lightness and Supp. Mat. Table S1a-b). Lightness or darkness are widely used as a proxy for melanisation (Gautam & Kunte, 2020; Hegna et al., 2013; Kutama et al., 2024; Lopez et al., 2021). GI, on the other hand, measures how green a pupae is (Yumnam et al., 2021). There was a strong positive correlation between lightness and GI across all experiments (Supp. Mat. Fig. S4). In nature, darker, more melanised, pupae are typically found on substrates away from leaves and on defoliated leaf midribs (off-leaf), while lighter, greener pupae are attached under leaves (on-leaf) (Clarke & Sheppard, 1972; Mayekar & Kodandaramaiah, 2017; Ohnishi, 1959; Sims, 1983; Yumnam et al., 2021). Results from Experiment 3 show that off and on-leaf substrates differ in their microclimate (Supp. Mat. Fig. S5). Off-leaf substrates are drier, being exposed to higher temperature and stronger winds. In contrast, on-leaf substrates are sheltered from direct sunlight, and transpiration from leaves may increase humidity and maintain cooler microhabitat (Deva et al., 2020; Lin et al., 2017). Therefore, the lighter, green pupae likely experience less desiccation stress than darker, melanised pupae. Consequently, larvae pupating under leaves may not need to allocate resources to physiological mechanisms for tolerating desiccation stress, unlike those pupating on off-leaf substrates. Under desiccation stress, greener pupae had poorer eclosion success compared to browner pupae supporting our prediction. However, neither lightness nor pupal colour was related to eclosion success when no desiccation stress was imposed (Experiment 5; Fig. 3 for lightness and Supp. Mat. Table S1b for GI). Together, these results support the idea that pupal colour plasticity is adaptive in the context of desiccation stress.

Melanin synthesis in insects is energetically costly (Ethier et al., 2015; Kutama et al., 2024; Roff & Fairbairn, 2013), and thus melanin should only be produced when it offers an adaptive advantage. In the tropical butterfly *Mycalesis mineus*, brown pupae were found more frequently under low RH conditions than under high RH (Mayekar & Kodandaramaiah, 2017), supporting the idea that melanisation has a role in desiccation tolerance. Similarly, studies in other insects demonstrate inter-population variation in melanin production wherein populations from arid environments tend to more melanised and more tolerant to desiccation than their counterparts in more humid conditions (Brisson et al., 2005; Parkash et al., 2009; Rajpurohit et al., 2008) These findings together support the idea that darker pigmentation provides a survival advantage in desiccating conditions either by enhancing water retention or mitigating the damage caused by stressors. In contrast, populations from consistently humid environments may not invest in melanin production, because the cost of synthesising melanin may outweigh its benefits under low desiccation stress (Britton & Davidowitz, 2025; Stoehr, 2006).

### No relationship between melanisation and water loss

One of the hypothesised mechanisms by which cuticular melanisation may prevent desiccation is the hydrophobic nature of cuticular melanin, which reduces cuticular transpiration (Hadley, 1994; King & Sinclair, 2015; Rajpurohit et al., 2008; Ramniwas et al., 2013). Some studies have indeed found a negative correlation between the degree of melanisation and water loss (King & Sinclair, 2015; Matute & Harris, 2013; Parkash, Sharma, et al., 2009; Ramniwas et al., 2013), while others have found no correlation (Britton & Davidowitz, 2024; Rajpurohit et al., 2016). In our study, we found no correlation between lightness of pupae and water loss. Interestingly, there was a negative correlation between lightness and eclosion success. This suggests that melanisation can enhance desiccation tolerance through factors other than reducing water loss. For example, a study on *Galleria mellonella* reported higher expression of stress management genes in melanic form than in non–melanic forms (Dubovskiy et al., 2013). Melanised individuals might also possess a greater capacity to heal the damages caused by diverse stressors or to prevent the damage from physiological stresses such as desiccation (reviewed in Dubovskiy et al., 2013; Solano, 2014). Melanin synthesis in insects affects multiple aspects of their biology (reviewed in San-Jose & Roulin, 2018). Thus, the processes and the products involved in melanin synthesis may confer a benefit under desiccating conditions even if melanin pigmentation does not directly limit cuticular water loss.

Most studies that tested the correlation between cuticular melanisation and water loss have been conducted on adult insects (Bujan et al., 2016; Parkash et al., 2009). Little is known about the structural and compositional properties of the pupal cuticle. The available evidence (e.g., Hopkins et al., 1984; Fraenkel & Rudall, 1947) suggests that cuticle properties – such as hardness, pigmentation, water content, and protein composition – change significantly across developmental stages. Given this, it is plausible that stage-specific differences in cuticular composition and organization could influence desiccation resistance differently. For instance, the presence, distribution, and layering of melanin in the adult cuticle may differ from those in the pupal cuticle, not only in quantity but also in how these melanins are structurally organized within the cuticular layers. Such differences in cuticle properties might influence cuticular permeability and water-retention properties differently in adults and pupae.

## SUMMARY AND CONCLUSIONS

We demonstrate that brown pupae have melanin in their cuticles. Under desiccating conditions, the melanised brown pupae, had a higher probability of eclosion than did lighter, green pupae. However, pupal colour did not provide any survival advantage under non-desiccating conditions. Our data show that desiccation stress is stronger on off-leaf microhabitats than under leaves. Because brown pupae tend to form on off-leaf substrates, pupal colour plasticity is likely an adaptation against desiccation. Water loss was inversely correlated with survival. Surprisingly, there was no correlation between melanisation and water loss, suggesting that pupal colour may confer tolerance to desiccation by mechanisms other than water loss, and highlighting the complexity of the relationship between melanisation, water retention and survival.

## Data Accessibility Statement

All data and code used for the analyses in this study will be archived in a publicly accessible GitHub repository, made available during peer review, and remain publicly accessible upon acceptance.

