## Supplementary file for "PUPAL COLOUR PLASTICITY AS A STRATEGY AGAINST DESICCATION"

### Supplementary file includes

**1) Text S1**: Quantification of colour.

**2) Text S2:** Melanin assay.

**3) Text S3:** Vapour Pressure Deficit (VPD) assay.

**4) Figure S1:** Pupation cylinders.

**5) Figure S2:** Histogram showing the frequency distribution of pupal GI for all experiments.

**6) Figure S3**: Frequency distribution of pupal greenness index (GI) for individual experiments.

**7) Figure S4:** Scatter plots showing the correlation between greenness index (GI) and lightness.

**8) Figure S5:**Line graphs showing differences in VPD between on-leaf and off-leaf substrates.

**9) Figure S6:**Raman spectra (1000 –1800 $cm^{-1}$) of brown and green pupal cases compared to a synthetic melanin standard.

**10) Table S1**: Model statistics of the Generalized Linear Models testing the effects of variables on eclosion probability.

#### Text S1: Quantification of colour

All pupae used in the analysis were photographed with a Nikon D3200 Digital SLR camera and a Sigma 105 mm macro lens. The following fixed settings were used: Aperture f/10, Exposure time 1/50th of a second, and ISO 800. A grey card (Model - JJC White Balance Gray Card Set GC-2) was used to correct white balance.

#### Text S2: Melanin assay

The assay was conducted on 8 green and 8 brown pupae. Lateral cuticular sections of dry pupal cases of green and brown pupae were fixed flat on glass slide using carbon tape on the side. Raman spectra were obtained using a Horiba Xplora Plus Confocal Micro Raman Microscope (Horiba Jobin, France). The samples were excited with a 532 nm at laser power settings of 0.1 and 10 mW, using 50× and 10× objective lenses, respectively. The accumulation times were 5 seconds for 10 mW and 3 seconds for 0.1 mW, and Raman shift values were collected between 1000 and 1800 $cm^{-1}$. At 0.1mW power, 12 pupal cases, six green and six brown, were scanned at two positions each, while at 10 mW, six pupal cases, three green and three brown were scanned at three positions each. Synthetic melanin (Sigma-Aldrich Chemical Corporation, M0418) was used as a reference. The reference sample underwent spectroscopic scanning without further processing.

#### Text S3: Vapour Pressure Deficit (VPD) assay

Dataloggers (Elitech, RC-4HC) were used to record temperature and RH from 08:30 to 17:00 h at substrates where green (on-leaf) and brown pupae (off-leaf) are typically found on *Cassia Fistula*, the host plant of *Eurema blanda* in the study locality. Data was recorded every minute, and for three days. Eighteen dataloggers in total were attached using tape under the midrib of intact leaves. Data from these dataloggers approximated the microclimate of green pupae. An additional eighteen dataloggers were attached to exposed branches or the midribs of defoliated leaves. Branches from nine trees were used for these microclimatic measurements, and these data approximated the microclimate of brown pupae. Vapour Pressure Deficit was calculated as $VPD = ((100 - RH)/100) * SVP$ where VPD is vapour pressure deficit in kPa, SVP is saturated vapour pressure and RH is relative humidity (Bujan et al., 2016; Law et al., 2020; Monteith & Unsworth, 2013).

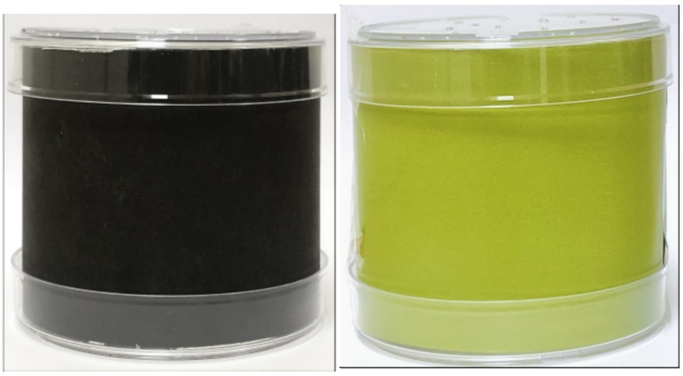

**Fig. S1: pupation cylinders.**

Examples of black and green paper cylinders used to obtain browner and greener pupae respectively.

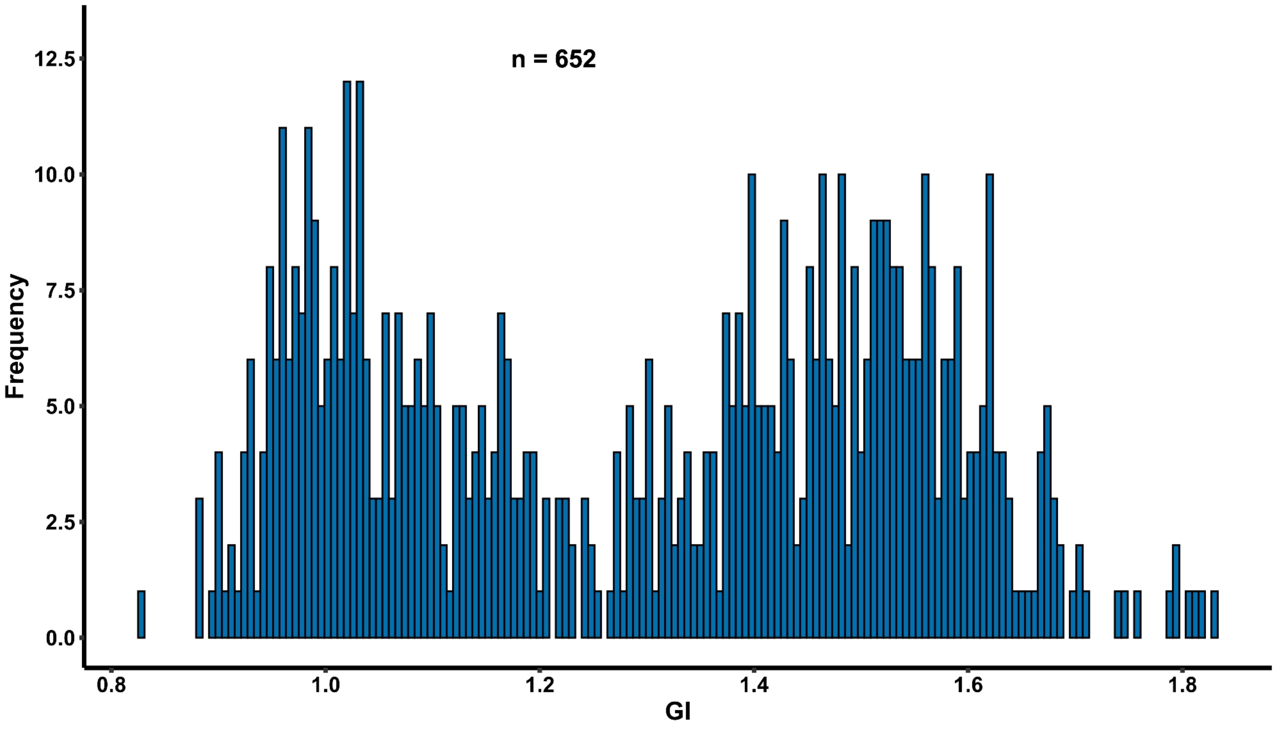


**Fig. S2: Histogram showing the frequency distribution of pupal GI.**

The X-axis represents the GI of pupae, and the Y-axis represents frequency. This figure includes data from all pupae from experiments 1, 4, 5, 6, and 7.

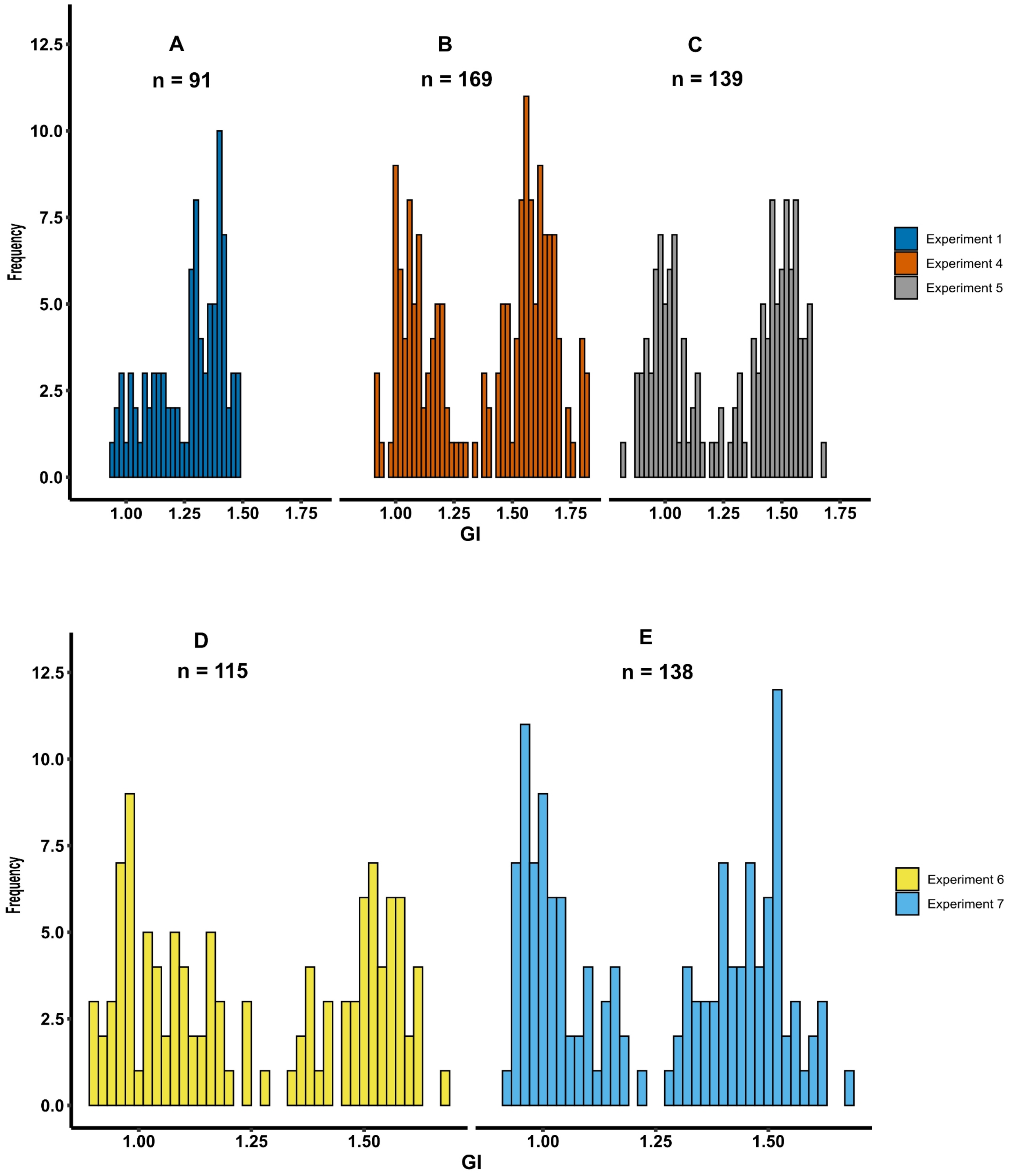


**Fig. S3. Frequency distribution of pupal greenness index (GI) for individual experiments.** The X-axis represents GI of pupae, and the Y-axis represents frequency. Panels: **(A)** Experiment 1, **(B)** Experiment 4, **(C)** Experiment 5, **(D)** Experiment 6, and **(E)** Experiment 7.

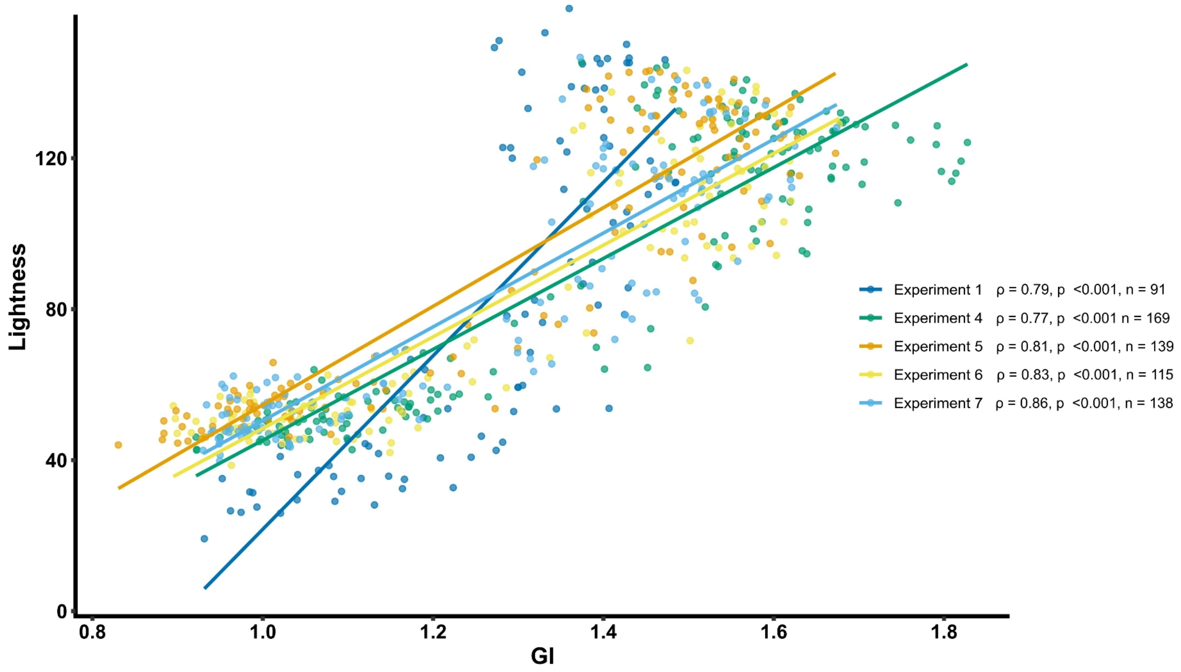


**Fig. S4. Scatter plots showing the correlation between greenness index (GI) and lightness.**

The x-axis represents GI, and the y-axis represents lightness. Regression lines and data points for experiments 1, 4, 5, 6, and 7 and depicted with distinct colours. ρ denotes Spearman's correlation coefficient, and *n* indicates the sample size per experiment.

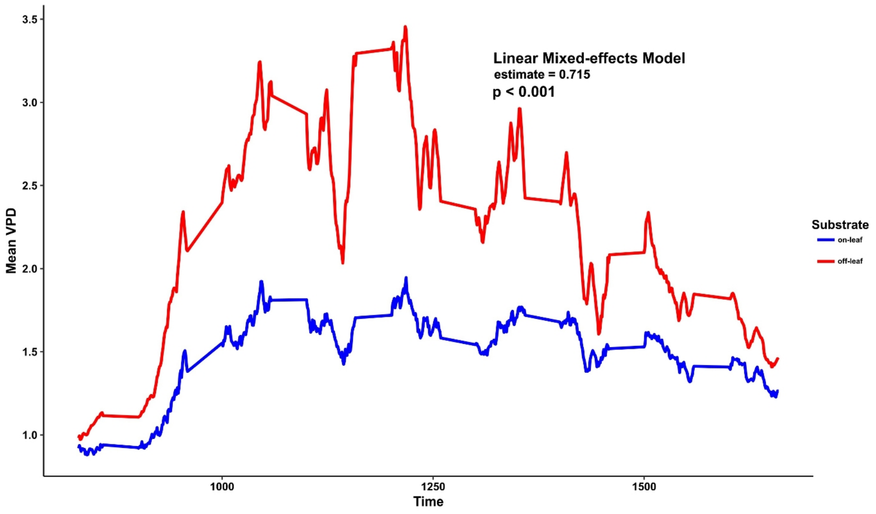


**Fig. S5: Line graphs showing differences in VPD between on-leaf and off-leaf substrates.**

The x-axis represents time of day, and the y-axis shows mean VPD (in kPa). The blue line represents on-leaf substrates, while the red line represents off-leaf substrates. Mean VPD differed significantly between the two substrate types.

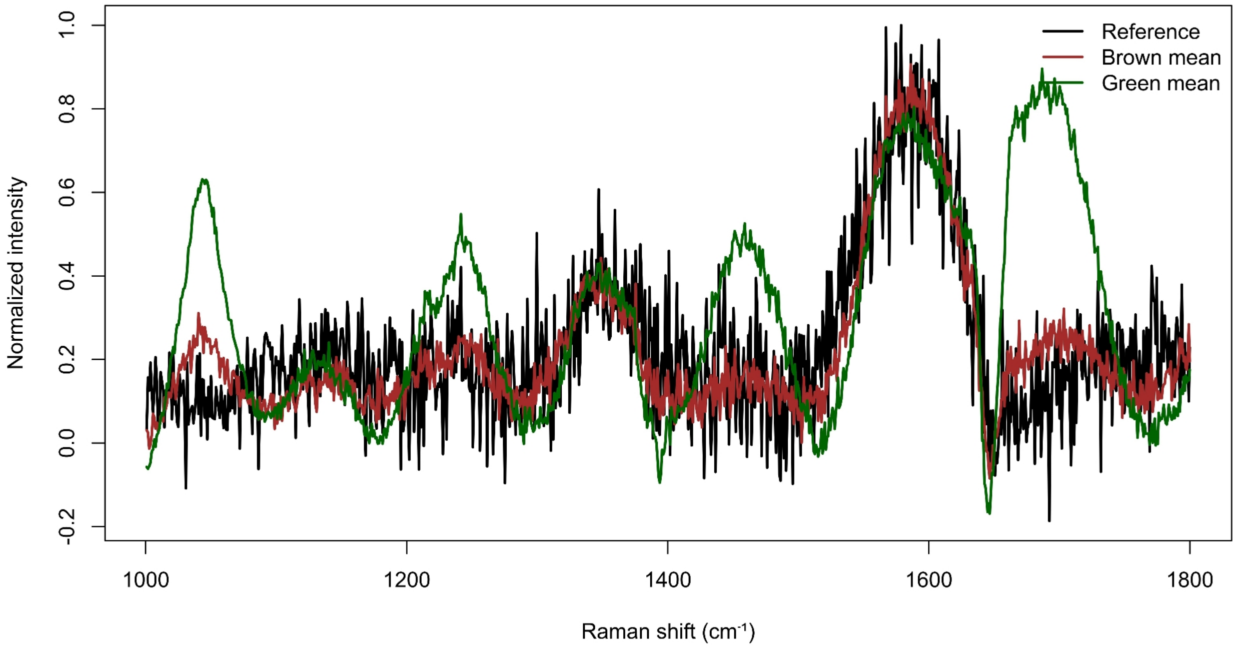


**Fig. S6. Raman spectra (1000 –1800 $cm^{-1}$) of brown and green pupal cases compared to a synthetic melanin standard.**

The pupal spectrum and reference spectrum was obtained at 10mW. All spectra were baseline corrected using asymmetric least squares and subsequently max-normalized. 3 brown and 3 green pupae were used to obtain the spectra. ˇFor each pupae two spectra were used for analysis. Spectra belonging to the same pupa were averaged to obtain a single mean spectrum. To generate group level mean spectra for green and brown pupal classes, individual spectra were averaged within each colour class (brown and green). The mean Raman spectra of brown and green pupae were then plotted against a single reference melanin spectrum over the 1000–1800 $cm^{-1}$. The x-axis represents Raman shift ($cm^{1}$), and the y-axis represents normalized intensity. The black line represents the synthetic melanin reference, the green and brown line represents mean spectra from 3 green, and 3 brown pupae respectively.

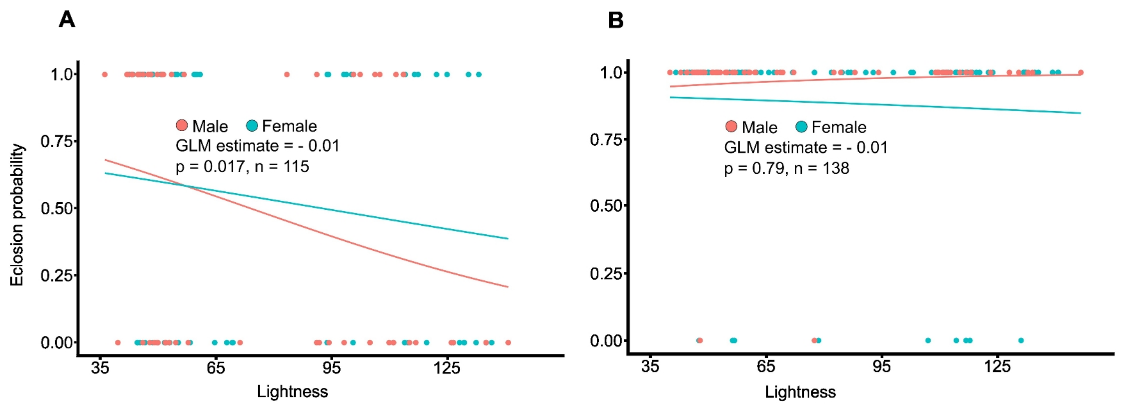


**Figure S7. Effect of pupal lightness on eclosion probability when larvae were reared and pupated at 60% RH: (A) under desiccation stress and (B) under non-desiccating conditions.** The dots represent data points, and regression curves for each sex are shown as solid lines. Estimates are for the independent effect of lightness on eclosion probability. The statistical results shown on the graph correspond to the combined data for males and females.

**.........................................................................................................................................**

#### Table S1: Model statistics of the Generalized Linear Models testing the effects of variables on eclosion probability

**Table S1a:** Comparison of additive and interaction models testing the effects of Lightness and Sex on survival with parameter estimates and AIC values

| Experiment and model formula | Variable | Estimate | Std. Error | z values | Pr(>\|z\|) | AIC |
|---|---|---|---|---|---|---|
| Experiment 4 Survival ~ Lightness +Sex | Lightness | -0.017 | 0.0058 | -2.929 | 0.003 | 174.08 |
| | Sex | -0.11 | 0.386 | -0.306 | 0.7596 | |
| Experiment 4 Survival ~ Lightness | Lightness | -0.019 | 0.008 | -2.258 | 0.023 | 175.95 |

| | | | | | | |
|---|---|---|---|---|---|---|
| *Sex | Sex | -0.560 | 1.299 | 0.431 | 0.66 | |
| | Lightness*Sex | 0.0042 | 0.011 | 0.357 | 0.72 | |
| Experiment 5 Survival ~ Lightness +Sex | Lightness | 1.77 | 1.99 | 0.892 | 0.372 | 9.55 |
| | Sex | 22.193 | 8846.5 | 0.003 | 0.998 | |
| Experiment 5 Survival ~ Lightness *Sex | Lightness | 1.77 | 1.99 | 0.89 | 0.372 | 11.56 |
| | Sex | 104.98 | 26720.4 | 0.004 | 0.997 | |
| | Lightness*Sex | -1.77 | 264.14 | -0.007 | 0.995 | |
| Experiment 6 Survival ~ Lightness +Sex | Lightness | -0.01 | 0.005 | -2.386 | 0.017 | 159.23 |
| | Sex | -0.21 | 0.384 | 0.547 | 0.58 | |
| Experiment 6 Survival ~ Lightness *Sex | Lightness | -0.009 | 0.007 | -1.245 | 0.213 | 160.45 |
| | Sex | 0.631 | 1.03 | 0.61 | 0.541 | |
| | Lightness*Sex | -0.010 | 0.012 | -0.87 | 0.380 | |
| Experiment 7 Survival ~ Lightness | Lightness | -0.002 | 0.009 | -0.261 | 0.793 | 78.315 |
| | Sex | 1.468 | 0.805 | 1.82 | 0.068 | |

| | | | | | | |
|---|---|---|---|---|---|---|
| +Sex | | | | | | |
| Experiment 7<br>Survival ~ Lightness*Sex *Sex | Lightness | -0.006 | 0.0108 | -0.61 | 0.53 | 79.459 |
| | Sex | -0.484 | 2.288 | -0.21 | 0.832 | |
| | Lightness*sex | 0.02 | 0.031 | 0.827 | 0.408 | |

**Table S1b:** Additive models testing the effects of GI and sex on survival with parameter estimates

| **Experiment** | **Variable** | **Estimate** | **Std. Error** | **z values** | **Pr(>\|z\|)** |
|---|---|---|---|---|---|
| Experiment 4 | GI | -2.46 | 0.814 | -3.03 | 0.002 |
| | Sex | 0.05 | 0.385 | 0.137 | 0.89 |
| Experiment 5 | GI | 24.4 | 23.07 | 1.058 | 0.29 |
| | Sex | 18.71 | 4473 | 0.004 | 0.99 |
| Experiment 6 | GI | -1.7 | 0.78 | -2.14 | 0.03 |
| | Sex | -0.13 | 0.38 | 0.34 | 0.73 |
| Experiment 7 | GI | -0.53 | 1.45 | -0.37 | 0.71 |
| | Sex | 1.45 | 0.804 | 1.813 | 0.07 |
